# Shared genomic signatures of Arctic adaptation among distantly related plant lineages

**DOI:** 10.64898/2026.10.26.762098

**Authors:** Cassandra Elphinstone, Marie Sandler, Philippa Stone, Maria Fernanda Torres Jiménez, Jonathan J. Henn, Xu Zhang, Siri Birkeland, Christian Brochmann, Greg H.R. Henry, Marco Todesco, Christine D. Bacon, Loren H. Rieseberg

## Abstract

Extreme environments impose strong selection pressures that can drive convergent adaptation among distantly related species. Arctic plants survive low temperatures, 24-hour light in summer and darkness in winter, short growing seasons, limited precipitation, nutrient-poor soils, and extreme seasonal variation. To test if these conditions are associated with shared genomic signatures, we generated a chromosome-level genome assembly for *Oxyria digyna* and compared it with existing reference genomes for three distantly related Arctic species (*Cochlearia groenlandica, Draba nivalis*, *Dryas octopetala*) and for 16 non-Arctic species from the same families. We identified 214 orthogroups with signatures of positive selection in all Arctic species, 1.6 times the number shared among the non-Arctic species. Although no gene ontology (GO) categories remained significantly enriched after multiple-testing correction, descriptive annotations included genes involved in general functions including protein folding, glycolysis, transcription, and oxidative-stress responses. Orthogroups showing copy number variation were associated with biotic interactions, sexual reproduction, and nutrient acquisition, suggesting that species may respond to biotic and nutrient stresses by shifts in gene copy number. Arctic and alpine genomes contained lower proportions of some repetitive elements, including terminal inverted repeats, and smaller regions of centromeric repeats than non-Arctic genomes. Overall, we see shared genes under positive selection, similar functions of genes with high copy number variation, and lower counts of some repetitive elements in the Arctic plant genomes. Our findings reveal a shared genomic signature of Arctic adaptation across deeply diverged plant lineages, suggesting that evolution in extreme environments repeatedly targets similar genomic features.

## Introduction

Natural selection often results in striking convergent morphologies and behaviors across distantly related species found in the same environment. Extreme environments such as deserts, alpine, and polar regions present natural laboratories for exploring the consequences of strong selection pressures on organisms and their genomes. The Arctic represents a dual challenge: it is among the most extreme environments on Earth and is currently experiencing one of the most rapid rates of warming (Rantanen et al. 2022). Plants have survived for millennia in the Arctic despite exposure to low temperatures, short growing seasons, relatively strong ultraviolet light, low-nutrient soils, and up to 24 hours of daylight in the summer (Cohen et al. 2014; Rantanen et al. 2022). Many Arctic plant species share traits that are clear adaptations to these extremes, including short stature, cushion growth form, dense pubescence, high root to shoot ratios, frost tolerance, loss of unnecessary pest resistance, low annual growth rates, and perennialism (Billings and Mooney 1968; Lütz 2010; Kevan and Godglick 2017; Bjorkman et al. 2018; Kemppinen et al. 2022; Gallois et al. 2025; Wang et al. 2025). With global warming forcing northward shifts in species ranges and reorganizing biotic communities, understanding how temperature adaptation interacts with Arctic light adaptations at a genomic level is essential (Wullschleger et al. 2015).

Recent advances in genomics show that the same genes, orthogroups/gene families, and/or gene networks are used repeatedly for adaptation, even in highly diverged species (Nocchi et al. 2024; Whiting et al. 2024), although one could expect that gene reuse should decrease with divergence time (Bohutínská and Peichel 2024; Satterlee et al. 2024). Chromosome-level genome assemblies enable us to detect convergent genomic changes in distantly related species that share an extreme environment, as is the case for Arctic species. This can help us better understand what parts of the genome are important for these adaptations and how repeatable evolution is at the genetic level (Zhang et al. 2023).

Many features of the genome, ranging from loci under positive selection to repetitive elements and gene locations, may be involved in adaptation and can be examined for signatures of convergent evolution. Molecular convergence does not necessarily occur only in homologous genes, but also in homologous genetic pathways through selection on different genes (Sackton and Clark 2019; Cerca 2023; Hoitinga and Birkeland 2025). In addition, orthogroup expansions and contractions are frequently consistent across species that share an environment (Zhang et al. 2023) and may reveal orthogroups that are functionally important or redundant, respectively.

While analyses of convergent evolution have often focused on coding regions, chromosome-scale genome assemblies make it possible to also identify patterns across the entire genome, including intergenic regions and variation in gene order (Wellenreuther et al. 2019). Genomic architecture is still relatively unexplored in the context of adaptation and convergence, and provides a broader context compared to analyses focusing only on gene-specific changes. This is particularly relevant because genes involved in adaptation to certain conditions are hypothesized to cluster in the genome, since genes near each other are often co-expressed (Williams and Bowles 2004; Zhao and Schranz 2019; Che et al. 2026) and strong linkage between locally adaptive genes may be favored by natural selection (Todesco et al. 2020; Battlay et al. 2023).

Short growing seasons and low temperatures are predicted to select for smaller genomes with fewer repeats, enabling faster cell divisions (Bureš et al. 2024). Because transposable element (TE) movement is also known to be affected by temperature and stress (Wos et al. 2021; Ito 2022), we hypothesize that TE repeat expansions and contractions may show convergence among Arctic species. While chromosome-level assemblies now allow us to compare repetitive elements between species, including transposable elements (Ito 2022), centromere types (Naish and Henderson 2024) and telomeric repeats (Fulnecková et al. 2013), the specific role of the environment in shaping centromeric and telomeric evolution remains largely unknown.

However, evidence that oxidative stress impacts telomere length (Barnes et al. 2019) and the discovery of unique telomeric repeats in some Arctic species (Kim et al. 2024) suggests that the environment may play a role in the evolution of these repetitive regions.

Previous comparative genetic studies of Arctic plants have focused on selfing Brassicaeae species (*Cardamine bellidifolia, Cochlearia groenlandica*, and *Draba nivalis*) and have primarily relied on transcriptomic data (Birkeland et al. 2020; Nowak et al. 2021; Birkeland et al. 2022). These studies revealed less convergence in genes under positive selection than might be expected based on comparative studies from other environments (Whiting et al. 2024), with shared evolutionary responses occurring primarily at the functional rather than the gene-specific level (Birkeland et al. 2020; Birkeland et al. 2022). Because these studies relied primarily on transcriptomic data, they were limited to capturing a snapshot of gene expression, potentially overlooking structural genomic changes, such as copy number variation and repeat expansions, that can drive long-term adaptation. Analyses of a chromosome-level genome assembly of *D. nivalis* showed that expanded orthogroups were associated with functions such as abiotic stress responses (e.g., oxidoreductase activity) and meiosis (Nowak et al. 2021), but without additional Arctic reference genomes it could not be assessed whether these patterns are shared across Arctic plant lineages. With the recent availability of chromosome-level reference genomes from more Arctic species (Billault-Penneteau et al. 2019; Kim et al. 2024), it is now possible to investigate whether similar genomic patterns underlie adaptation across independent Arctic lineages. Notably, in the extreme environments of the Arctic, selfing mating systems and small effective population sizes may strengthen drift, reduce the efficacy of selection, and constrain adaptive responses.

Here we present a new chromosome-level genome assembly for the outcrossing Arctic species *Oxyria digyna* and compare it with existing reference genomes for three distantly related Arctic species (the two selfing species *Draba nivalis* and *Cochlearia groenlandica* and the outcrossing species *Dryas octopetala*) and for 16 non-Arctic species from the same families.

This dataset provides a rare opportunity to test for genomic convergence across lineages separated by up to ∼125 million years of evolution (Kumar et al. 2017). We hypothesize that (1) Arctic lineages will share more signatures of positive selection and recurrent orthogroup expansions and contractions relative to non-Arctic lineages; (2) Arctic species will have lower TE abundances and shorter putative pericentromeric regions than their non-Arctic relatives; and (3) Arctic species will share microsyntenic blocks that are not found in closely related non-Arctic relatives.

By integrating chromosome-level assemblies from multiple Arctic and non-Arctic lineages, this study tests whether adaptation to the Arctic produces repeatable signatures across genes, orthogroups, repetitive elements, and genome architecture. This study provides a foundation for predicting how Arctic plant lineages may respond genomically to the ongoing rapid environmental change in the Arctic. Beyond the Arctic, this study addresses a fundamental question in evolutionary biology: to what extent does the environment repeatedly affect genomic architecture across large phylogenetic distances?

### Study species

The four target Arctic vascular plant species were the only ones with reference genomes at the time of study (Supplementary materials A). We excluded species from the Arctic category that mainly occur in alpine areas and are absent from the High Arctic (e.g. *Arabis alpina*, *Thlaspi arvense*, and *Rheum nobile* (Koch et al. 2006; Geng et al. 2021; Hu et al. 2022; Feng et al. 2023). These alpine species were compared as a separate category in some analyses in this study (e.g. comparing repeat content between the alpine, Arctic and non-Arctic-alpine species). The four target Arctic species have circumpolar ranges including the most extreme northern areas of the High Arctic. However, only two of the reference individuals were sampled in the High Arctic: *Oxyria digyna* from Alexandra Fiord on Ellesmere Island (79°N) (Supplementary materials B) and *Cochlearia groenlandica* from Spitsbergen, Svalbard (79°N) (Kim et al. 2024). The *Dryas octopetala* reference plant was grown outside Edinburgh, Scotland for several years after having been collected from West Sutherland in the far north of mainland Scotland (58°N) (Darwin Tree of Life), and the *Draba nivalis* reference plant was collected from the southern edge of the Alaska Range (∼62°N) (Grundt et al. 2006; Nowak et al. 2021).

The outcrossing, perennial herb *Oxyria digyna* (Polygonaceae) is diploid (2*n* = 14) throughout the Arctic and has a genome size of approximately 600 Mbp estimated by kmer analysis (Kim et al. 2024) and flow cytometry (protocol in the Supplementary materials C); hexaploids (2*n* = 42) have only been found in the Himalayas (Aiken et al. 2007). The long-lived (up to 100 years) and outcrossing dwarf shrub *Dryas octopetala* (Rosaceae; diploid with 2*n* = 18) has a genome of 229 Mbp. The diploids *Draba nivalis* (2*n* = 16) and *Cochlearia groenlandica* (2*n* = 14) belong to the Brassicaceae and are predominantly selfing, perennial herbs, and have genome sizes of 302 Mbp (Nowak et al. 2021) and 250 Mbp, respectively (Kim et al. 2024). Our non-Arctic species contained both perennials and annuals, and the shrubs/trees in Rosaceae and the herbs in Polygonaceae are typically longer-lived than the herbs in Brassicaceae.

## Methods

We built a chromosome-level genome assembly for diploid *O. digyna* and compared it to the genome assemblies of three other diploid Arctic species (*D. nivalis, C. groenlandica,* and *D. octopetala,*) and 16 non-Arctic species from the same families (Table 1; diverged ∼90-130 Mya a TimeTree plot can be found in the Supplementary materials B): *Fagopyrum esculentum, F. tataricum, Polygonum aviculare, Rheum tanguticum,* and *R. nobile,* (Polygonaceae); *Arabidopsis lyrata, Arabis alpina, Brassica oleracea, Capsella rubella,* and *Thlaspi arvense* (Brassicaceae); and *Argentina anserina, Fragaria vesca, Malus sylvestris, Prunus persica, Pyrus bretschneideri,* and *Rosa rugosa* (Rosaceae) (Magallón et al. 2015; Kumar et al. 2017; Zhang et al. 2017). We compared 1) orthogroups with signals of positive, relaxed and intensified selection; 2) orthogroup expansions and contractions; 3) transposable element repeat types and abundance; 4) centromere types and sizes; 5) regions with shared gene order (microsynteny); and 6) gene locations relative to centromeres. Code used for the analyses is available at https://github.com/celphin/Oxyria_genome.

### Gene annotations

We ran a modified version of a plant-specific MAKER annotation Snakemake pipeline (Lazaro 2023). The pipeline uses MAKER version 3.01.03 (Campbell et al. 2014) and RepeatMasker 4.1.5 (Tarailo-Graovac and Chen 2009) with the RMBlast search engine to mask repeated sequences. We created an annotation of haplotype 1 of our *Oxyria digyna* assembly (detailed description of the assembly methods can be found in the Supplementary materials B), as well as of the larger haplotype of the *Dryas octopetala, Fagopyrum tataricum,* and *Polygonum aviculare* assemblies. Peptide sequences were determined with the ORFik library (Tjeldnes et al. 2021). To improve the annotation, we ran the pipeline using an annotation of *Rheum tanguticum* for protein homology evidence in the species from the Polygonaceae. The gff3 files were converted to cds files using bedtools 2.30.0 (Quinlan and Hall 2010). BUSCO v5.1.2 (Manni et al. 2021) was run against the eudicots_odb10 database to assess the quality and completeness of the protein annotation. Available gene annotations were downloaded from NCBI.

For the Arctic species, only annotations with Annotation Edit Distance (AED) scores of 0.6 were used for further analyses. The functions of the resulting protein sequences were determined with InterProScan v.5.64-96.0 (Quevillon et al. 2005; Jones et al. 2014). We used the following databases (defaults for InterProScan) to classify proteins: TIGERFAM, SFLD, SUPERFAMILY, PANTHER, Gene3d, Hamap, ProSiteProfiles, Coils, SMART, CDD, PRINTS, PIRSR, ProSitePatterns, AntiFam, Pfam, MobiDBLite, PIRSF, with the settings: ‘*Gene Ontology lookup’* enabled and ‘*precalculated lookup for identical protein matches’* disabled (Ashburner et al. 2000).

We used ErmineJ 3.2 (Gillis et al. 2010) to run gene ontology (GO) enrichment analyses to determine the enrichment of putative functions of orthogroups identified in several of the following analyses. Default settings were used with significant enrichment defined at a *p-value* threshold of 0.01, the *‘bigger is better’* option for count scores and the *–log-transform* option for *p-value* scores; analyses were run for all three types of GO terms (biological process, cellular component, and molecular function).

### Orthogroup comparisons

We ran OrthoFinder 3.0 (Emms and Kelly 2019) once with default settings on all species from the three plant families (in total 20 species). Files were parsed for OrthoFinder using GENESPACE to enable downstream analyses of synteny. STAG (Emms and Kelly 2018) and STRIDE (Emms and Kelly 2017) were used to reconstruct the most parsimonious and rooted species tree from the many gene trees in OrthoFinder. The resulting dichotomously branching, rooted species tree was made ultrametric using the *chronos* function in the ape package in R, which estimates divergence times using a penalized maximum likelihood method (Paradis and Schliep 2019). We supplied an approximate root divergence time range for the three angiosperm families of between 90-130 Mya, based on estimates from (Magallón et al. 2015; Kumar et al. 2017; Zhang et al. 2017).

To test for positive selection on the Arctic branches in each orthogroup, we used the adaptive branch-site random effects likelihood method (aBSREL) implemented in HyPhy v2.5.49 (Smith et al. 2015). To prepare the data from OrthoFinder for HyPhy, we aligned the proteins within each orthogroup to each other using the phylogeny-aware aligner PRANK (Löytynoja 2014). As protein alignment errors are known to increase the rate of false positives (Jordan and Goldman 2012; Privman et al. 2012), we filtered our alignments with GUIDANCE2 (Landan and Graur 2008; Sela et al. 2015) using 10 bootstrap replicates. Columns with low confidence scores (<0.9) and sequences with low scores (<0.9) were removed. The filtered aligned protein sequences were translated back into codon sequences using PAL2NAL (Suyama et al. 2006) and the original coding sequences. Terminal branches corresponding to the four Arctic species were designated a priori as foreground branches on the gene trees inferred by OrthoFinder, and aBSREL was run on each orthogroup to test for episodic positive selection.

Within the HyPhy software we used RELAX to test for relaxed or intensified selection on the Arctic branches (Wertheim et al. 2015). Significance was assessed by BH-FDR across orthogroups; k > 1.2 indicates intensification and k < 0.8 indicates relaxation. P-values were corrected for multiple testing within each orthogroup tested with HyPhy, but not between tests as orthogroups should be independent with regard to selection (Smith et al. 2015). Although trees containing paralogs do not violate the assumptions of HyPhy, paralogs may confound cross-lineage interpretation through orthology errors, gene conversion and heterogeneous constraints. Therefore, we compared orthogroups containing paralogous copies with those containing single-copy genes in the Arctic species to minimize potential confounding effects of paralogs (Casola and Hahn 2009).

To test whether genes found to be consistently under positive selection in the Arctic species were also under positive selection in the non-Arctic species, we checked counts and overlap between orthogroups under positive selection in the four most closely related non-Arctic species. Gene ontology enrichment analyses for the functions of the Arctic branches under positive selection were run in ErmineJ as described above. We used a background gene set for each species based on all genes tested in aBSREL with gene ontology annotations.

To assess orthogroup enrichment, we used CAFE5 (Mendes et al. 2021). We used the dichotomously branching, rooted, ultrametric tree and the counts of the phylogenetic hierarchical orthogroups built in OrthoFinder for all species. We excluded orthogroups that were missing in one or more species because this can result in falsely inflated orthogroup expansions and contractions. CAFE5 was run using the *-e error model* option 50 times. Orthogroups that were identified as expanded, contracted, or rapidly evolving in 47/50 of the runs (94% of the time) were kept and considered ‘real’ expansions, contractions, and significant rapid changes. CAFEplotter (https://github.com/moshi4/CafePlotter) was used to graphically display the results.

### Centromere comparisons

Putative centromere locations were predicted using RepeatOBserverV1 for all species with chromosome-level assemblies (we did not have a chromosome-level genome assembly for *Cochlearia groenlandica* at the time of analyses) (Elphinstone et al. 2025). Putative centromere types were classified from the RepeatOBserver heatmaps as retrotransposon-based, tandem repeat-based, or mixed. Putative retrotransposon-based centromere locations were predicted at genome positions with the maximum repeat summed abundance on each chromosome, while putative tandem repeat-based centromere locations were predicted as minimums in the repeat Shannon diversity index for each chromosome (Elphinstone et al. 2025).

### Gene duplication types

For the three Arctic species with chromosome-level assemblies, we used DupGen finder (Qiao et al. 2019) to search for and classify duplicated gene pairs resulting from whole-genome duplication (WGD), tandem duplication, proximal duplication, transposed duplication or dispersed duplication. Whole genome duplication genes were identified by their appearance in syntenic blocks shared between chromosomes. We ran DupGen finder on *Oxyria digyna* (with *Polygunum aviculare* as outgroup), *Dryas octopetala* (with *Rosa rugosa*) and *Draba nivalis* (with *Capsella rubella*) with the default parameters using the gene BLAST results from OrthoFinder as input.

For each Arctic species, we defined core and accessory genes using the OrthoFinder results. Core genes were defined as genes that appeared in all 20 species and accessory genes as genes that only appeared in the species of interest (possibly resulting from TE duplications). For each Arctic species, we plotted the fraction of core and accessory genes relative to the total number of genes in 2 Mbp windows across the genome.

### Transposable element frequencies

Transposable elements were annotated with the Extensive de-novo TE Annotator (EDTA) v2.0.1 using default settings (––sensitive 0 ––anno 1 ––evaluate 0 ––threads 40 ––force 1) (Ou et al. 2019). Summaries of the abundance of each TE family in each genome (Arctic and non-Arctic) were compiled from the EDTA.TEanno.sum files. Transposable element positions from the EDTA.TEanno.gff3 files for *O. digyna* and *D. octopetala* were plotted along the genome using ggplot2 in R (v4.4.0) (Wickham 2009).

### Microsynteny between Arctic genomes

We used GENESPACE 1.4 (Lovell et al. 2022) to compare synteny among all the species in each family as well as among the three Arctic species with chromosome-level assemblies (*Draba nivalis, Dryas octopetala,* and *Oxyria digyna*). Orthologs were identified by OrthoFinder. Syntenic blocks were grouped by MCScanX (Wang et al. 2012) and plotted by GENESPACE. To identify micro-syntenic regions in the Arctic species that are not found in their non-Arctic relatives, we extracted the orthogroups shared among all species. For each species, we calculated the minimum number of genes found between genes associated with each pair of orthogroups. We extracted all the orthogroup pairs that had nearby genes in all the Arctic species (within 40 genes of each other) and were found farther apart in the genomes of the non-Arctic species (more than 40 genes apart). This pairwise comparison followed the same steps as the SynNet pipeline (Zhao and Schranz 2019) but did not build a network of orthogroups; rather it compared all possible pairs of orthogroups. We also determined gene ontology and InterProScan terms for genes consistently found near each other only in Arctic species.

### Gene location comparisons

Each gene type (positively selected, in microsynteny clusters, core and accessory) was counted in 2 Mbp windows along the genome, and the relative gene density was calculated as a fraction of the total gene count in that window. Each window was assigned as either centromere-distal or centromere-proximal based on its proximity to the putative centromere, with the closest half of the chromosome defined as proximal and the rest as distal. A two-sided Wilcoxon test in R was used to determine if there were significantly different relative abundances of each gene type in centromere-distal versus centromere-proximal regions. To determine if there were more positively selected genes than expected in the microsyntenic regions, we compared the observed overlapping counts to the counts we would expect by random chance (p-value <0.05).

## Results

### Genome assembly and annotations

Our assembly of *Oxyria digyna* had a haploid genome size of 582 Mbp, an N50 of 74.7 Mbp, an assembly BUSCO of 93.4%, an annotation BUSCO of 83.7%, and > 90% of the assembly was included in the seven chromosomes in both haplotypes (**Error! Reference source not found.**). The annotations for *Dryas octopetala, Oxyria digyna,* and *Polygonum aviculare* consistently had BUSCO scores 5-10% lower than the assembly BUSCO score (Table 0).

### Orthogroup comparisons

Using OrthoFinder, we analyzed 634 489 genes from the 20 species (Table 1) and placed 608 021 of them into 33 157 orthogroups, of which 8 168 were species-specific and 7 588 shared by all species. From the OrthoFinder results, we built a rooted, dichotomously branching, ultrametric species tree (Figure 1A). We inferred that *Oxyria* diverged from *Rheum* ∼39 Mya, and that *Dryas* diverged from the *Malus/Prunus* clade ∼41 Mya, aligning with previous studies (Magallón et al. 2015; Xiang et al. 2017). TimeTree analyses (Supplementary materials D) were also consistent with our species tree from OrthoFinder and provided a maximum divergence time of 125 Mya (Kumar et al. 2017).

**Figure 1:**
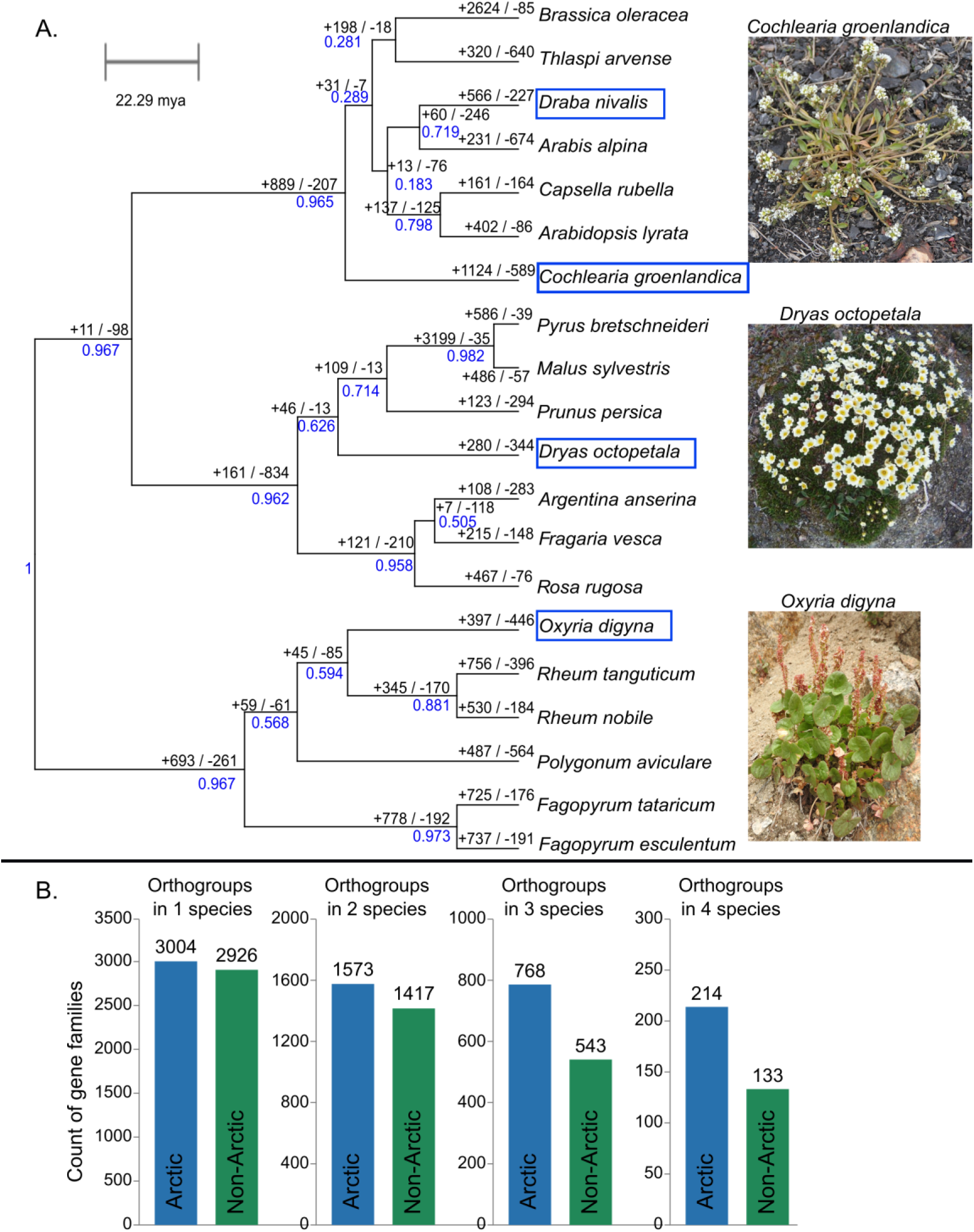
(A) Rooted, binary, ultrametric species tree from OrthoFinder and ape chronos showing CAFE expanded (+) and contracted (−) orthogroup counts shown in black. The branch support is shown in blue. Photos of an Arctic species from each family are included. Photos of *Oxyria digyna* and *Dryas octopetala* were taken by Cassandra Elphinstone. The photo of *Cochlearia groenlandica* is by Bjoertvedt (CC BY-SA 3.0, https://commons.wikimedia.org/w/index.php?curid=22417146) (B) Comparing counts of orthogroups showing signatures of positive selection in Arctic species versus non-Arctic species. Fewer orthogroups were found showing signatures of positive selection in the four non-Arctic species compared to the overlap in all four Arctic species.

Due to high divergence levels, only 10 493 orthogroups contained genes that could be successfully aligned to each other and converted back into the coding sequence. We detected branch-site evidence of positive selection in all four Arctic species for 214 orthogroups (Table 2). We compared these results with the amount of overlap in four of the most closely related non-Arctic species, some of which are alpine species, (*Arabidopsis lyrata, Arabis alpina, Prunus persica,* and *Rheum nobile*) and identified 133 orthogroups under positive selection across the four non-Arctic species (Figure 1B). Of the 214 orthogroups consistently showing positive selection in all four Arctic species, sixteen orthogroups were also under positive selection in the four most closely related non-Arctic species. Using a permutation test, we randomly selected four species from the combined Arctic and non-Arctic species pool. At least 214 orthogroups showed evidence of positive selection across all four species in about 1.5% of random trials (p = 0.015; 5,000 permutations). At least 133 orthogroups showed evidence of positive selection across all four non-Arctic species in about 2.6% of trials. These permutation results suggest that the observed overlap is greater than expected under this random-selection model, including among non-Arctic species. After excluding paralogs, we found signatures of positive selection in 122 orthogroups shared across all Arctic species and in 87 orthogroups in all four non-Arctic species (Table 2). The positively selected genes shared by all four Arctic species did not show significant gene ontology enrichment, and nearly all of them fell into broad functional categories related to protein folding, membrane lipids, DNA repair, and oxidative stress.

Among the 8 613 orthogroups shared by at least two Arctic species, we detected significantly intensified selection (k > 1.2) in 62 and significantly relaxed selection (k < 0.8) in 52. Only one orthogroup, associated with carbon-carbon lyase activity (GO:0016830), showed significantly intensified selection in all four Arctic species (Table 2). Two orthogroups showed significantly relaxed selection in three of the four Arctic species, one was associated with protein folding, and one had an unknown function.

No orthogroups were consistently expanded (had multiple paralogous copies) in all four Arctic species, but 29 orthogroups (including two with rapid expansion; significantly higher rate of orthogroup expansion than across the rest of the tree (p < 0.05)) were expanded in three species. These included orthogroups with potential growth regulation functions, including *Tryptophan Aminotransferase-Related 1 (TAR1)* involved in auxin biosynthesis and *MAINTENANCE OF MERISTEMS*). Three contracted orthogroups were found in all four Arctic species with InterProScan functions related to ATP binding sites in two of them and apoptosis in the third.

We observed far greater functional convergence (i.e. orthogroups belonging to the same functional GO category) in orthogroup expansion and contraction among all four Arctic species than at the level of specific orthogroups, for which convergence was low. Expanding and contracting orthogroups shared several of the same functions, so we could only link many functions to a change in size but not specifically expansion or contraction (Table 3; Supplementary material E). In addition, some functions were associated with both expansion and contraction of orthogroups in the same species. For example, orthogroups associated with xenobiotic transport were both expanded and contracted in *Cochlearia groenlandica, Dryas octopetala,* and *Oxyria digyna* (Table 3).

### Centromere comparisons

There were smaller regions of pericentromeric repeats in the three Arctic species’ chromosomes than in their non-Arctic relatives (Figure 2). However, we found no consistent putative centromere types in the Arctic genomes (Table 4). Two of them had putative locations composed of retrotransposon centromeres; whereas the third had mostly imperfect tandem repeats (two example chromosomes are shown in Figure 2). The centromeric repeats in *Oxyria digyna* and *Dryas octopetala* resembled those of their non-Arctic relatives, whereas *Draba nivalis* had long terminal repeat (LTR) retrotransposon-based centromeres. Putative centromeres composed of nearly perfect tandem repeats were found in *Polygonum aviculare*, and similar, but more mutated tandem repeats were found in *Oxyria digyna* (Figure 2) and *Rheum nobile* (Table 4). *Fagopyrum* spp. had a mixture of retrotransposons and tandem repeats. Putative retrotransposon centromeres were found in several species of the Rosaceae, including *Dryas octopetala* (Figure 2), *Malus sylvestris*, *Rosa rugosa,* and *Rubus idaeus*, and putative tandem repeat centromeres were found in *Argentina anserina* and *Fragaria vesca* (Table). In contrast to *D. nivalis*, most other Brassicaceae species had clear tandem repeat-based centromeres (e.g. *Arabidopsis thaliana*, *Arabidopsis lyrata*, *Brassica rapa*).

**Figure 2:**
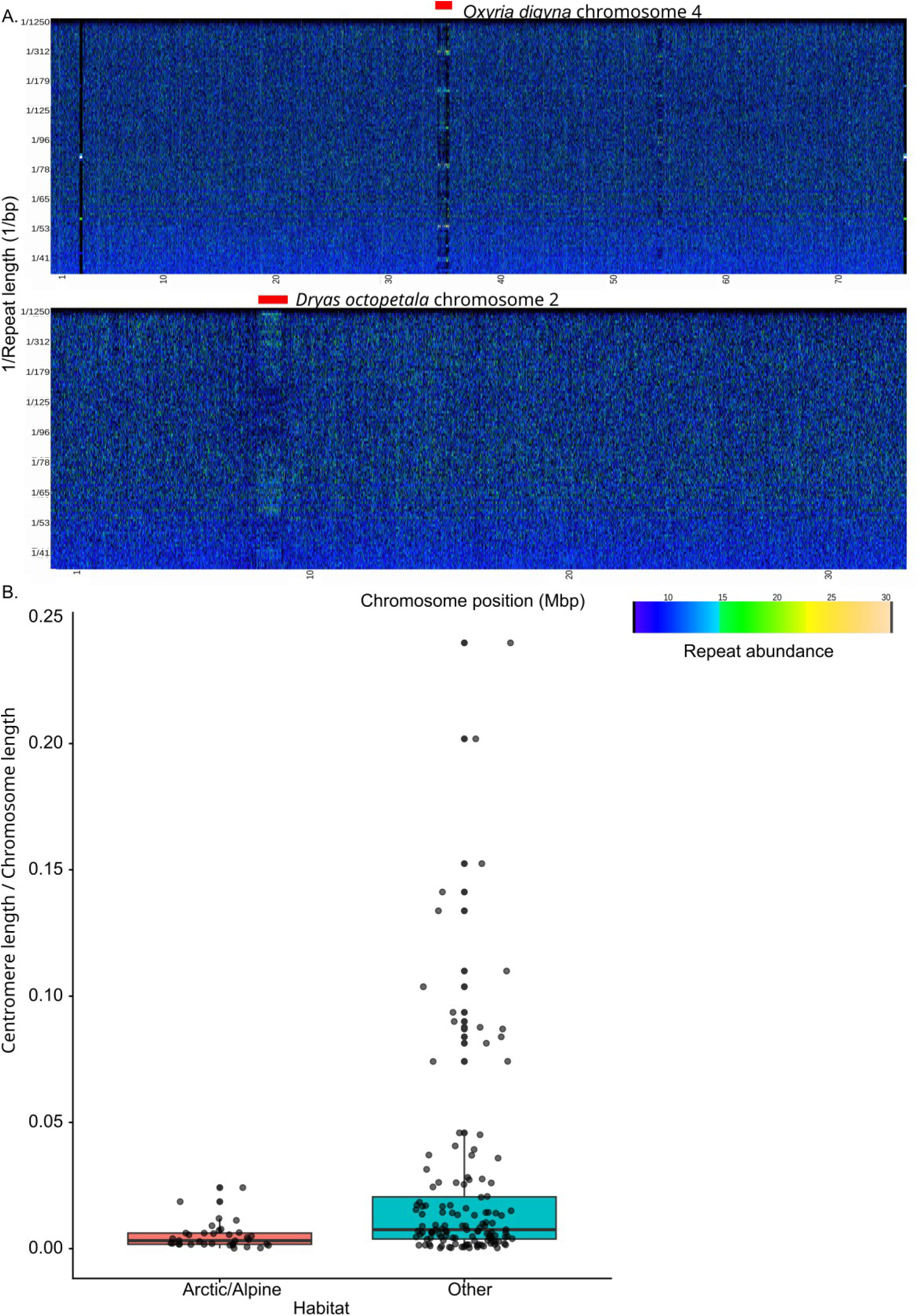
(A) RepeatOBserver spectra plots for *Oxyria digyna* chromosome 4 (top) and *Dryas octopetala* chromosome 2 (bottom). These examples show a retrotransposon centromere in *D. octopetala* and a tandem repeat centromere in *O. digyna*. The red rectangles indicate the putative centromere location. (B) Boxplot showing the relative centromere lengths for the Arctic/alpine species compared to the other species. Each dot represents a chromosome.

### Transposable elements

Overall repeat abundances were not lower in the Arctic species than the non-Arctic ones (Table 5). However, the combined group of the Arctic and the non-Arctic but alpine species (e.g. *Arabis alpina* and *Thlaspi arvense*) showed significantly lower abundances of terminal inverted repeats than the other non-Arctic species (*p* = 6.19 e-06, specifically hAT (Kruskal-Wallis *p* = 0.027), CACTA (*p* = 0.044), Tc1_Mariner (*p* = 0.044); Figure 3). As expected, Gypsy long terminal repeat retrotransposons were abundant at the putative centromere locations in *Dryas octopetala, Oxyria digyna,* and *Draba nivalis*.

**Figure 3:**
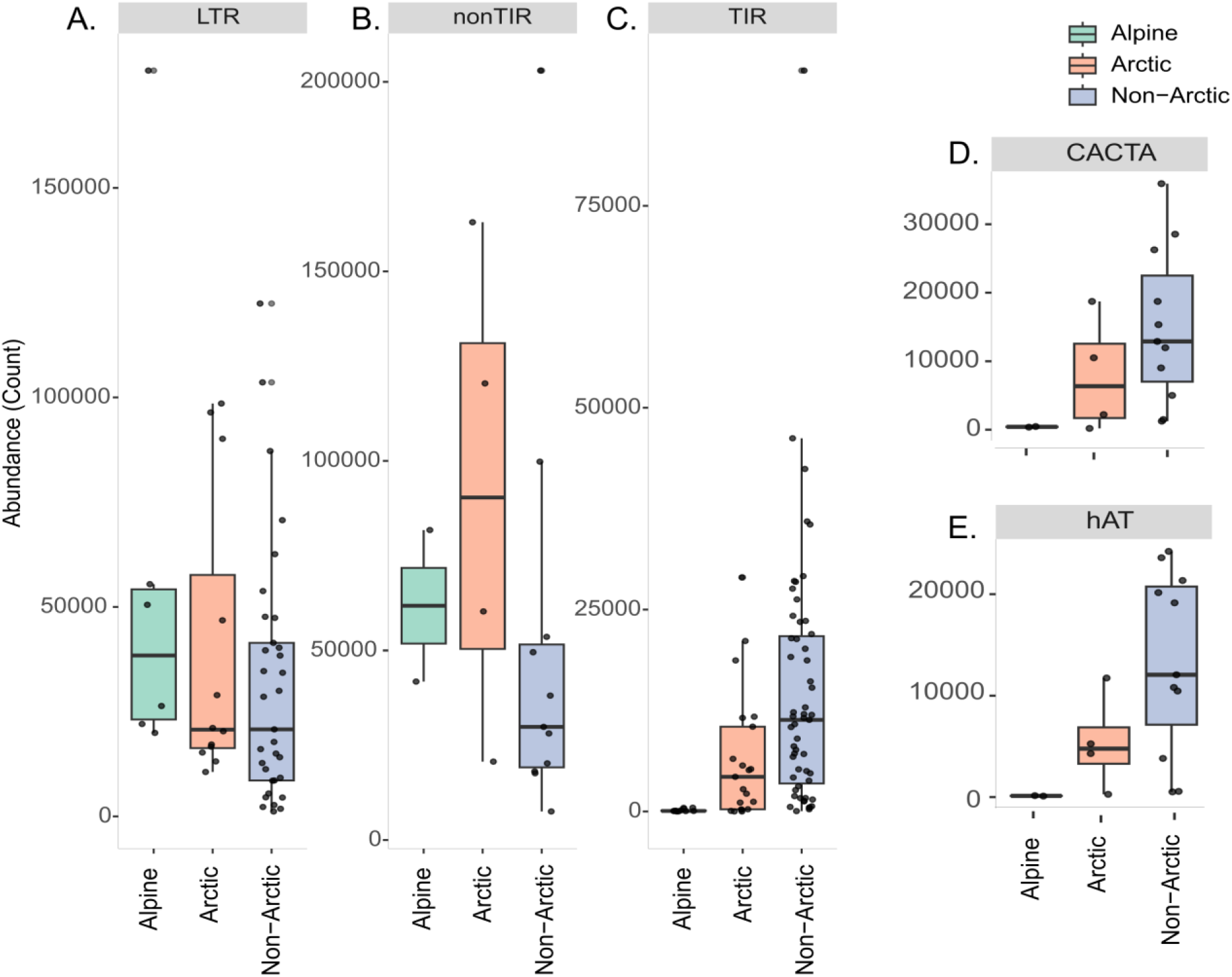
Transposable element repeat abundances (A, B, C) in Arctic (*Draba nivalis, Dryas octopetala, Oxyria digyna,* and *Cochlearia groenlandica*), alpine (*Arabis alpina* and *Thlaspi arvense*), and non-Arctic-alpine species. Arctic and alpine species have less terminal inverted repeats (TIRs) than non-Arctic species (Kruskal-Wallis test; *p* = 6.19 e-06). Specific TIRs driving this pattern are the hAT (D) and CACTA (E) repeats.

### Gene duplications

Dispersed duplicate genes, replicated genes found dispersed throughout the genome with no ancestral syntenic pattern, were the most common type of gene pairs in all species (Table 6). Of the Arctic species, *Dryas octopetala* had the largest number of gene duplication pairs (64 732) of which 28% (18 345) were whole genome duplication (WGD) pairs, duplicated groups of genes found with the same genomic synteny across different chromosomes. These were consistently found at high frequency near the putative centromere locations and may therefore not be caused by real WGD. Most (91%) of the centromere-associated WGD gene pairs in this species were found in all chromosomes, implying syntenic centromeres. In *D. nivalis* and *O. digyna*, only a few (11% and 17%, respectively) of the gene duplication pairs were associated with WGDs. The WGD pairs in *O. digyna* were found in all chromosomes with varying densities and were not associated with the putative centromere locations.

*Oxyria digyna* had the second largest number (56 525) of gene duplication pairs of the four Arctic species, of which 16 104 (28%) were transposed duplication pairs. In both *Dryas octopetala* and *O. digyna,* we found tandem and proximal duplicated genes were much more abundant near the telomeres, far from the centromeres, likely due to higher total gene densities in these regions. In *Dryas octopetala*, these duplicated gene pairs were associated with GO terms involved in environmental stress responses, such as 84 paralogs of Cytochrome P450 (UV and drought responses), 35 paralogs of UDP-glycosyltransferase (cell wall modifications and plant development), and 20 paralogs of small auxin-up RNA (development and growth regulation). All categories of duplicated gene pairs (whole genome, tandem, transposed, proximal, dispersed) peaked near the putative centromere locations in *Dryas octopetala* but not in the other species.

### Microsynteny between Arctic genomes

The three Arctic species with chromosome level assemblies showed some regions of shared synteny (microsynteny) even though they diverged 90-125 Mya (*Draba nivalis, Dryas octopetala,* and *Oxyria digyna*; Table 7, Figure 4A). We assessed gene order patterns shared between these three Arctic species and between these Arctic species and their non-Arctic relatives (*Arabidopsis lyrata, Prunus persica,* and *Rheum nobile*). We find that the amount of shared microsynteny was not specific to the Arctic. We found many gene pairs that were consistently separated by less than 40 genes uniquely in all three Arctic species, but there were also multiple long clusters of such genes in both the Arctic-Arctic and Arctic-non-Arctic comparisons. For example, we observed 16 clusters shared between *Dryas octopetala* and *Oxyria digyna* and 24 shared between the *Dryas octopetala* and the non-Arctic *Polygonum aviculare*.

**Figure 4:**
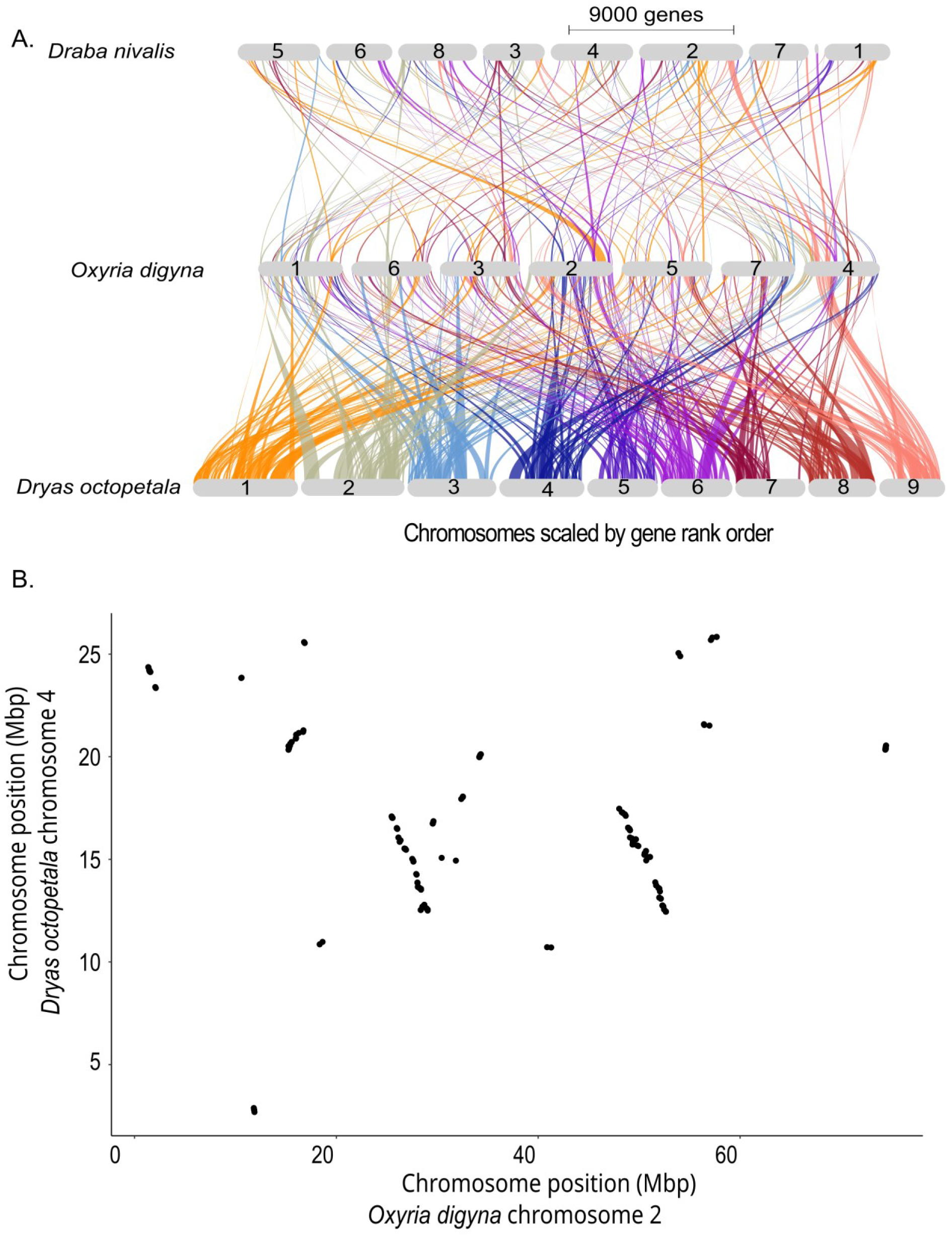
Microsynteny between Arctic genomes. (A) GENESPACE synteny plot of Arctic species. Chromosomes (grey ovals) are ordered based on synteny with the *Dryas octopetala* genome. Syntenic regions are coloured based on the *Dryas* chromosomes. (B) An example large, shared gene cluster between 12 and 18 Mbp on chromosome 4 in *Dryas octopetala* corresponds to two clusters at different locations in *Oxyria digyna* (Chr 2: 28-30 Mbp and 50-52 Mbp). This set of genes are consistently found near each other in all three Arctic plant genomes (*Draba nivalis, Dryas octopetala* and *Oxyria digyna*).

The Arctic-specific shared gene clusters (genes found within the 40 gene distance in all three Arctic species and not in the non-Arctic species) often contained genes with functional annotations associated with homeobox, MADS-box domains, or transcription factors (Table 7; Figure 4). For example, one Arctic-specific cluster included a homeobox domain, multiple copies of *HEAT INTOLERANT 4*, sugar transporters, and a programmed cell death protein. The microsynteny regions shared between the Arctic species overlapped multiple times with the centromeric regions in *Oxyria digyna* but never in *Dryas octopetala* (Figure 5).

**Figure 5:**
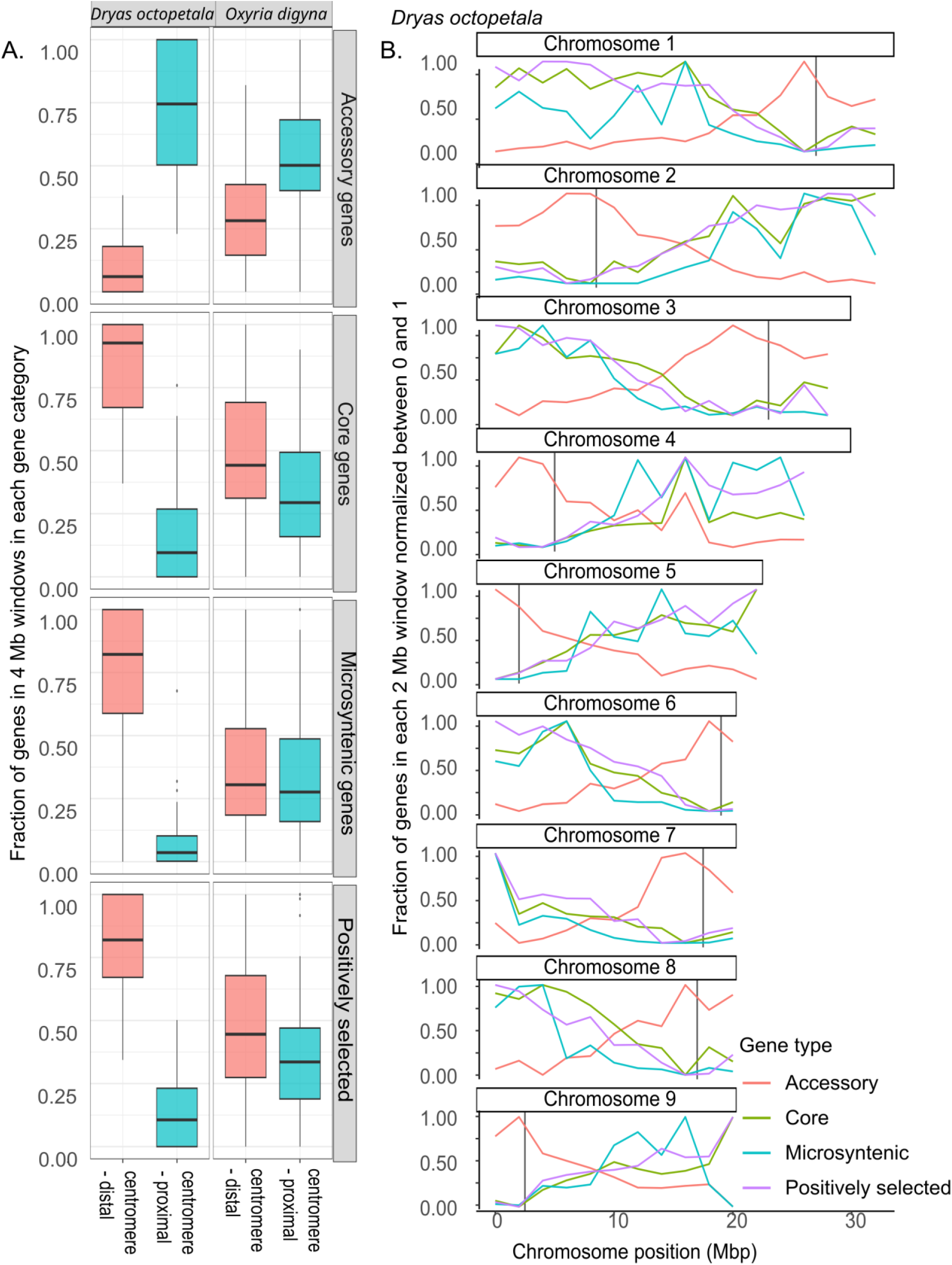
(A) Gene abundances relative to the centromere position. Accessory genes are most frequent centromere-proximal regions, whereas core genes, genes in microsyntenic regions, and positively selected genes are most frequent in regions farther from the centromeres (centromere-distal). (B) Abundance of gene types plotted along the *Dryas octopetala* chromosomes. Vertical lines represent the centromeres.

### Genomic locations of positively selected genes, microsyntenic regions, and core genes

Positively selected genes, microsyntenic regions, and core genes were significantly enriched far from the centromeres (p-values in Table 8), while LTR repeats, duplicated genes, and accessory genes showed enrichment around the centromeres. The pattern of core gene abundance near the telomeres and accessory gene abundance near the centromeres was in all 20 species. In the Arctic species, this pattern was most clear in *Dryas octopetala*, which had high abundance of retrotransposons at the centromeres and was telocentric, but it was also apparent in *Draba nivalis* and *Oxyria digyna* (Figure 5). We observed 788 positively selected genes in the microsyntenic clusters, but such overlap is expected 93% of the time for two sets of randomly selected genes in the *Dryas octopetala* genome of the same size as the number of positively selected genes.

## Discussion

What genomic characteristics do Arctic plants share? Our comparative genomics study of distantly related Arctic plant genomes identified similarities in branch-site tests for positive selection, centromere sizes, and repeat composition, even across 125 Mya of evolutionary divergence. The genomic similarities across Arctic species suggest these characteristics may have evolved repeatedly in response to the Arctic environment.

Nearly 1.6x the number of orthogroups showed signatures of positive selection in all four Arctic species compared to the overlap between their non-Arctic relatives. This was a clear excess of Arctic convergent selection beyond the non-Arctic background, concentrated in a small set of genes. The excess is only evident when genes are shared across species. For example, single-species counts are almost identical (3,004 Arctic vs. 2,926 non-Arctic), and then the excess overlap in the Arctic grows with the number of species sharing them (+11%, +41%, +61%). The positively selected orthogroups in the Arctic species had surprisingly broad, housekeeping gene ontology terms. Although housekeeping genes are not expected to be under directional selection, similar gene categories have been found to be under positive selection in divergent species adapting to different climates (Whiting et al. 2024) and alpine ecosystems (Rellstab et al. 2020; Zhang et al. 2023). While we did not observe expansions or contractions in the same orthogroups across Arctic species, there was an enrichment in orthogroups belonging to the same GO functional categories (Table 9), suggesting that there are some processes that are important for Arctic adaptation, even if their regulation is not achieved through the same genes in different Arctic species. Overall, we see different scales of genomic convergence from signatures of positive selection occurring in the same orthogroups across highly diverged species, the same functional categories for orthogroup expansion and contractions, and specific repeat types contracting in the Arctic environment.

### Orthogroup convergence

The number of orthogroups that showed significant signatures of positive selection in the branch-site tests was surprisingly high, given earlier work on three Arctic Brassicaceae found limited overlap (Birkeland et al. 2020). However, this previous study was based on transcriptomes and used a deliberately conservative approach (one sequence per species from each orthogroup, strict alignment filtering, branch-site tests per clade). Our approach tested entire gene families from whole genome data and is therefore more sensitive, while the comparison with non-Arctic relatives partly controls for background convergence. Our findings are also consistent with those of several studies on alpine environments, where multiple positively selected, expanded, and contracted orthogroups were identified (Rellstab et al. 2020; Zhang et al. 2023). In these studies, the convergent orthogroups were associated with similar GO functions as those we observed in the Arctic species studied here, including DNA repair, lipid membrane composition, reproduction, and stress resistance (Rellstab et al. 2020; Zhang et al. 2023).

The orthogroups under positive selection included many genes involved in core functions like transcription and protein folding. This suggests that genes involved in basic cell processes frequently show signatures of positive selection in many species, which aligns with the types of genes previously observed to be involved in local adaptation in very divergent species (Whiting et al. 2024). However, we notably observed nearly double the number of shared genes under positive selection among the Arctic species, suggesting that shared environmental conditions create a higher level of convergence at the gene level.

### Repeats and centromeres

We identified some terminal inverted repeats (TIRs; CACTA and hAT transposable elements) as well as the centromeric repeats, that are at much lower abundances in the Arctic and alpine plants. The relatively small regions of pericentromeric repeats in all three Arctic species may be linked to assembly quality versus true centromere lengths, but no assembly metrics indicated this was the cause. Overall, we did not find fewer repeats in the Arctic/alpine species, as might be expected in lower quality assemblies. These normal levels of total repeats was contrary to the prediction that the short growing seasons and cold temperatures in both Arctic and alpine environments should select for smaller genomes with fewer repeats (Bureš et al. 2024). Both CACTA and hAT TEs, like many TEs, are well known for changing gene expression and triggering chromosomal rearrangements, often creating phenotypic variation that selection can act on (Belyayev et al. 2022). CACTA TE families have been explored in numerous other species for their adaptive insertions into various genes (McClintock 1950; Belyayev et al. 2022). Consistent with our results, CACTA and hAT TEs were less expressed in high than in low-elevation ecotypes of *Arabidopsis arenosa* (Wos et al. 2021). However, we are not aware of other studies reporting reduced abundance of these TEs families in Arctic or alpine species. An exception to this pattern was *Draba nivalis,* which has normal TIR abundances, suggesting that surveys of additional species will be needed to determine the strength of this apparent pattern.

The Arctic plant centromeric repeat types were most similar to their non-Arctic relatives with the exception of *Draba nivalis*, in line with previous findings that centromere composition and repeat structure are partially consistent with species phylogenies (Henderson et al. 2025).

### Gene locations

We hypothesized that genes important for adaptation to Arctic environments would be preferentially retained in microsyntenic blocks or conserved genomic neighborhoods. Consistent with this hypothesis, we identified clusters enriched for homeobox genes and other transcription factors, suggesting that selection may preserve regulatory neighborhoods involved in developmental and environmental responses (Irimia et al. 2012) in all three chromosome-level Arctic genomes and not in their closest relatives. Interestingly, these microsyntenic regions, along with positively selected genes and core genes, clustered away from the centromeres while accessory genes (those unique to only one species), duplicated genes, and many repeat classes were concentrated in the centromere-proximal regions. This pattern can be explained by differences in recombination rates in different regions of the chromosome, with high recombination rates away from the centromeres increasing the efficacy of selection. However, we did not see significant overlap between the microsyntenic genes and the positively selected genes, as we might expect in regions uniquely important for adaptation to Arctic environments (such as response to 24 hours of light and short growing seasons). Some of these conserved microsyntenic regions may function as integrated regulatory modules that coordinate developmental timing, flowering, and environmental response, enabling Arctic plants to synchronize growth and reproduction within extremely narrow and variable seasonal windows. The conserved synteny around *Hox* genes observed in studies of metazoans may be due to regulation by cis-acting elements (Irimia et al. 2012), possibly making any recombination near them deleterious. Surprisingly, we found that in the *Dryas octopetala* genome but not the *Oxyria digyna* genome, these clusters were enriched in higher recombination regions away from the putative centromeres.

Across all the species (Arctic and non-Arctic) with chromosome-level assemblies that we studied, we observed that core genes were typically centromere-distal, whereas accessory genes were enriched in centromere-proximal regions. Because centromeric regions experience reduced recombination (Zhang et al. 2026), this suggests that many accessory genes are more tightly linked compared to core genes.

In *Dryas octopetala* more than 2,000 genes were annotated as whole-genome-duplication-derived genes, and nearly all were located near putative centromeric regions. Many of these genes were associated with GO terms related to retrotransposon-like functions, including DNA integration and reverse transcriptase activity (Neumann et al. 2011; Xin et al. 2024). The relative scarcity of WGD genes elsewhere in the genome suggests that these duplications do not reflect a recent whole-genome duplication event but instead likely represent localized accumulation of transposon-derived sequences within centromeric regions. These transposon-derived sequences may also be linked to the formation of some accessory/de novo genes.

We did not observe a higher abundance of transposon genes near the *D. nivalis* and *O. digyna* centromeres, possibly because the centromeric regions are enriched in tandem repeats and not retrotransposons. However, in these species we still see an abundance of both transposons and accessory genes near the centromeres on the majority of chromosomes. Our finding of more tandem and proximal duplicated genes near the telomeres is consistent with higher recombination rates typically observed near telomeres (Zhang et al. 2026) and the likely origin of tandem duplicated genes from unequal crossing over (Silver 2001). Tandem and proximal gene duplicates have been shown to display similar gene expression patterns (Qiao et al. 2019).

Together, these patterns suggest that different genomic regions may play distinct roles in gene duplication and adaptation, with centromeric regions potentially acting as reservoirs for new accessory genes and telomeric regions containing core genes under the strongest purifying selection.

### Haplotype variation

Genomes of individuals can vary a lot within species, such as in copy number and structural variations, which can facilitate local adaptation (Lowry and Willis 2010; Todesco et al. 2020; Hämälä et al. 2024). The *Dryas octopetala* and *Draba nivalis* individuals used in this study (had the only complete chromosome level genomes) were collected below the Arctic Circle and could differ significantly from other individuals of the same species found above the Arctic Circle, especially when considering the Arctic was recolonized by a subset of individuals from lower-latitude refugia that could survive at (and adapt to) higher latitudes (Elphinstone et al. 2024; Brochmann et al. 2025). More pangenomes for Arctic/alpine species are needed to explore this intraspecific variation.

## Conclusions

Divergence time matters when testing for evolutionary repeatability since the probability of gene reuse in repeated adaptation is expected to decline as lineages diverge (Bohutínská et al. 2021; Bohutínská and Peichel 2024). Given that the Arctic species included in this study diverged up to 125 million years ago, the overlap we identify is notable. We observed convergence of specific orthogroups under positive selection and convergence of functional categories repeatedly expanding or contracting in all studied Arctic species. Our findings suggest that adaptation to the Arctic environment may repeatedly involve similar genes and pathways, but many of these genes are also often under selection in other non-Arctic species as well. Knowing the amount of molecular convergence between Arctic plant genomes will reveal how repeatable evolution in the Arctic is and help predict how non-Arctic species might adapt to Arctic conditions as they track their temperature niches polewards.

## Data availability

The *Oxyria digyna* genome assembly and associated sequencing data are available in the NCBI BioProject database under accession numbers PRJNA1175703 and PRJNA1175704. Analysis scripts and related files are available at https://github.com/celphin/Oxyria_genome

## Acknowledgments

An NSERC Vanier, a Weston doctoral scholarship, and an NSERC PDF to CE enabled this project. The Digital Research Alliance of Canada was used for all high-performance computations. Sequencing and CDB were supported by the Swedish government-funded Strategic Research Area Biodiversity and Ecosystem Services in a Changing Climate, BECC and the Swedish Research Council (#2017-04980; #2022-03927). Seed was collected on a fieldwork trip to Ellesmere Island in 2018 funded by an NSERC Discovery grant and logistical support from the Polar Continental Shelf Programme to GHRH. A research permit to sample plants was obtained from the Nunavut Department of Environment in 2018.

## Author contributions

Cassandra Elphinstone: field *Oxyria digyna* seed collections, grew reference *O. digyna* plant, *O. digyna* flow cytometry, *O. digyna* HMW extractions, *O. digyna* genome assembly bioinformatics, genome annotations of *Dryas* species and *O. digyna* genomes, ran comparative genomics analyses of all Arctic and non-Arctic species, wrote manuscript and made revisions. Marie Sandler: genome annotations for multiple Polygonaceae species, assisted with comparative genomics analyses of all species Arctic and non-Arctic species, and suggestions on manuscript. Marco Todesco: HiC library preparations and lab work guidance, writing/editing of all manuscript drafts. Siri Birkeland: conceptual analysis design, guidance on alignment and positive-selection analyses, interpretation of results, writing/editing of manuscript drafts. Christian Brochmann: writing – review and editing. Greg Henry: Ellesmere field sampling funding, edited manuscript drafts. Philippa Stone: *Oxyria digyna* karyotyping and recommendations on manuscript. Maria Fernanda Torres Jiménez: attempted initial DNA extractions, helped obtain sequencing funding, and worked on figure design. Jonathan Henn: helped obtain sequencing funding. Loren Rieseberg: Funded lab work, conceptual analysis design, writing/editing of manuscript drafts. Christine D. Bacon: Funded HMW and HiC sequencing, edited manuscript drafts.

## Tables

**Table 1: Genomes compared in this paper, with genome assembly and annotation statistics, OrthoFinder and CAFE results.**

**Table 2: Counts of genes families showing positive selection in aBSREL and RELAX in HyPhy comparing the overlap between four Arctic species and four non-Arctic close relatives. Arctic species show nearly double the overlap compared to non-Arctic species.**

**Table 3: Expanded and contracted orthogroups gene ontology (GO) terms. Amino acid related terms are highlighted in yellow, sugar/glycosylation in pink, oxidative stress in cyan, sexual reproduction in red, nutrient and vitamins in orange, methylation in purple, DNA damage and repair in grey, cell wall or microtubule related in light blue, responses to abiotic stimulus (light, temperature, water) and biotic interactions in green, and tetrapyrrole related in dark green.**

**Table 4: Centromere types for Arctic species and their close non-Arctic relatives.**

**Table 5: Transposable element repeat abundances for each species described in Table 1. Annotations of TEs were made with EDTA.**

**Table 6: Gene duplication counts (whole genome (WGD), tandem (TD), proximal (PD), transposed (TRD), and dispersed (DSD) duplications) for the Arctic-alpine species *Oxyria digyna, Draba nivalis, Dryas octopetala* and *Arabis alpina*.**

**Table 7: Microsynteny: Gene pairs found within 40 genes of each other in *Draba nivalis, Dryas octopetala* and *Oxyria digyna*. InterProScan gene functional descriptions are coloured by frequently found categories in these regions: “conserved site” – pink, “EF-hand” –blue, “homeobox” – green, “transcription factor” – purple, and “active site” – orange**.

**Table 8: Gene enrichment in centromere-proximal versus centromere-distal regions of the genome. Each gene type was counted in 2Mbp windows along the genome, and the relative gene density was calculated as a fraction of the total gene count in that window. Each window was assigned as either centromere-proximal or centromere-distal based on its proximity to the putative centromere with the closest half defined as proximal and the rest as distal. A two sided Wilcox test in R was used to determine if there were significantly different relative abundances of each gene type in centromere-distal versus centromere-proximal regions of the genome. P-values and q-values are listed for each gene category.**

**Table 9: Arctic environmental pressures and possibly associated gene ontology (GO) terms. DN – *Draba nivalis*, DO – *Dryas octopetala*, CG – *Cochlearia groenlandica*, OD – *Oxyria digyna*, all – all four Arctic species**.

## Notes

https://github.com/celphin/Oxyria_genome

### Competing Interest Statement

The authors have declared no competing interest.

