## Supplementary_materials for "Shared genomic signatures of Arctic adaptation among distantly related plant lineages"

### Supplementary material

A: Study species

The focal Arctic species in this manuscript occur in the most extreme high latitudes, which are defined as zones A-C (the Arctic polar desert to the Arctic shrub tundra) in the Pan-Arctic Flora (Elvebakk, 1999; Walker et al., 2005).

*Oxyria digyna* (Mountain Sorrel, 2n = 2x = 14, 600 Mbp, Polygonaceae)

The genus *Oxyria* appears to have diverged from *Rheum* 15-40 mya (Zhang et al., 2022). The only other species in the genus is *Oxyria sinensis,* which is an alloploid derived from *O. digyna* and a ghost lineage (Luo et al., 2017). *Oxyria digyna* seems to have originated on the Tibetan Plateau and migrated into the Russian Arctic about 11 mya, from where it moved eastwards into North America and westwards into Europe (Wang et al., 2016). The current circumpolar populations of *O. digyna* are thought to have spread out of the Himalayas about 3 mya (Allen et al., 2012). The Polygonaceae also contains the genus *Rumex* with multiple dioecious species (Grant et al., 2022), has low synteny between species compared to the Brassicaceae and has multiple species that are allopolyploids (Kim et al., 2008). *Oxyria digyna* has been reported as predominantly dioecious (having separate male and female plants), but bisexual (hermaphrodite) plants have been reported in some populations (Brochmann & Steen, 1999). The species is thus predominantly outcrossing, but hermaphrodites can also self. *Oxyria digyna* is a long-living perennial and important to the Inuit and Sami Peoples for its high vitamin C content and use as reindeer fodder (Aiken et al., 2007). This species shows tremendous plasticity in response to temperature and drought, and numerous ecological studies have focused on this species to study flowering time responses to temperature (Bjorkman et al., 2015) and local adaptation (Bjorkman et al., 2016).

*Dryas octopetala* (Mountain Avens, 2n = 2x = 18, 260 Mbp, Rosaceae)

There are four *Dryas* species (*D. integrifolia, D. alaskensis, D. ajenesis, D. drummondii*) with contig-level reference genomes (Billault-Penneteau et al., 2019; White & Pirro, 2022). *Dryas octopetala* also has a complete chromosome-level assembly (Darwin Tree of Life, NCBI Genbank ID GCA_963921425.1). The genus was recently proposed as a model system for root symbiosis (Billault-Penneteau et al., 2019) because *D. drummondii* (and some *D. integrifolia* (Henry & Svoboda, 1986) , but not *Dryas octopetala*) can fix nitrogen through symbiotic relationships with *Frankia* and other rhizobia bacteria in root nodules. *Dryas* species also form ectomycorrhizal relationships with a few fungal species to enable colonization of glacial forelands (Gardes & Dahlberg, 1996). Many species of the Rosaceae, in contrast to the Polygonaceae, have similar base chromosome numbers (x = 7-9) and synteny is typically high (Davik et al., 2022).

*Draba nivalis* (Snow Whitlowgrass, 302 Mbp, 2n = 2x = 16) and *Cochlearia groenlandica* (Polar Scurvygrass 250 Mbp, 2n = 2x = 14) (Brassicaceae)

The Brassicaceae is one of the most thoroughly studied plant families partly because many of the species in this genus have small diploid genomes and can easily be compared to the model plant *Arabidopsis thaliana,* also found in the Brassicaceae.

B: *Oxyria digyna* genome assembly

Seeds were collected in August 2019 at Alexandra Fiord, Nunavut, Canada (78.9112° N, 76.1455° W) and stored in coin envelops at -20 ºC. The reference sample germinated in June 2021 (Figure 2) on filter paper with 24-hour light at 21 ºC. After about 14 days, germinated seedlings were moved to a mix of peat and perlite. The reference plant flowered after about four months in a growth chamber at 16-19 ºC and 24 hr light. All the following protocols were conducted on this one individual approximately one year after germination. For chromosome counting, some rosettes were broken off, and clones were successfully grown in hydroponics (using a mixture of ¼ Hoagland solution and tap water) to provide clean root tissue.

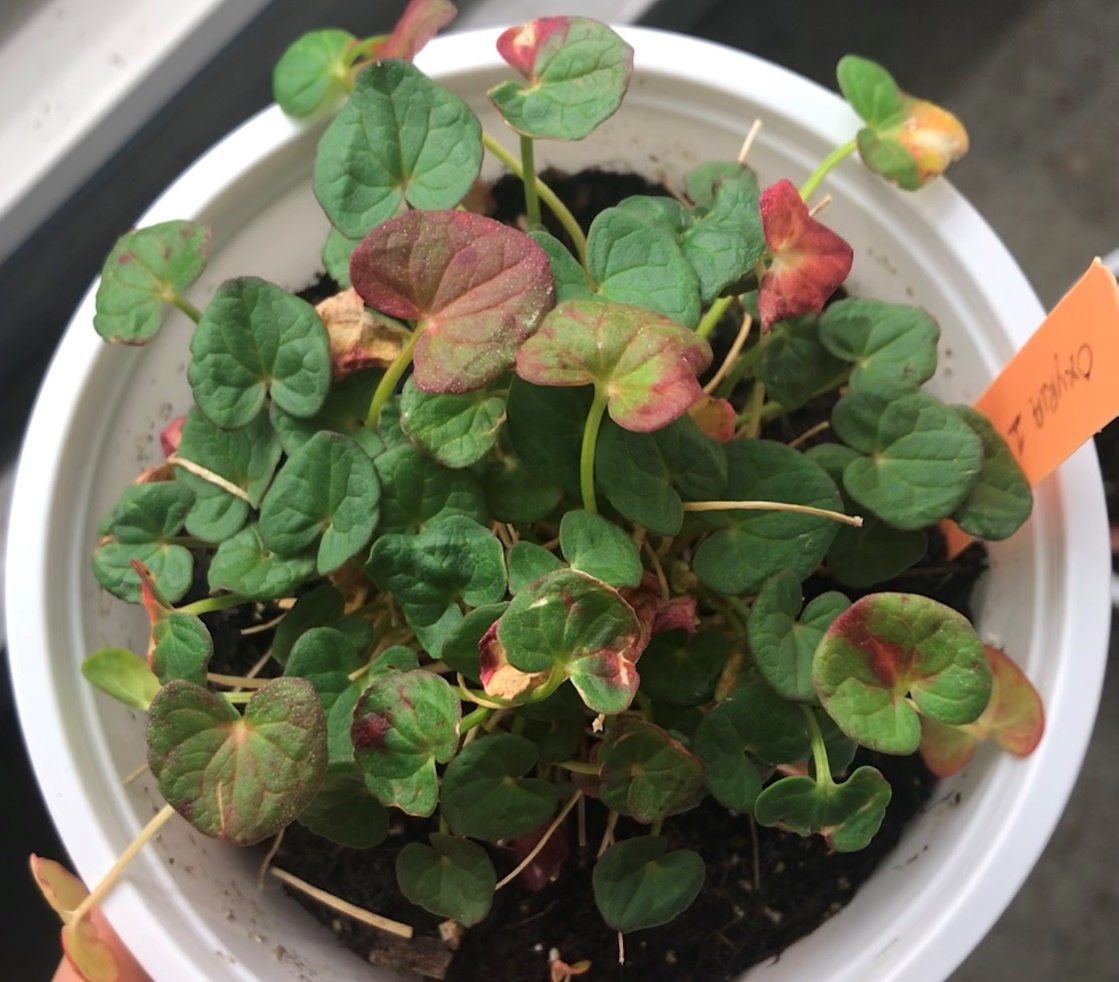

Figure 2: The reference *Oxyria digyna* plant grown from seed collected at Alexandra Fiord, Ellesmere Island, Nunavut, Canada. Photo credit Cassandra Elphinstone.

To determine the chromosome number, we used the Feulgen squash technique with cold hydrolysis following (Jong, 1997) and Schiff’s reagent (Fox, 1969) for staining. The plant was kept at 4 ℃ for 24 hours prior to chromosome squashes. We fixed, hydrolysed, and stained cells from the root apical meristems, and viewed them under 40x magnification. We verified that the reference individual was diploid with 14 chromosomes based on previous precise chromosome counts (Farooq & Saggoo, 2014) and not hexaploid as reported in some populations in the Himalayas.

To estimate the genome size of *O. digyna* we performed flow cytometry with fresh leaf tissue and the standard *Solanum lycopersicum* (2C = 1.96 pg DNA). This standard was selected because it has a genome size that is between 2x and 0.5x that of many Polygonaceae species (Kew Gardens C-values database). We used a Cytoflex flow cytometer in the Biomedical Research Center at the University of British Columbia and followed the modified protocol described in (Supplementary materials C ).

Fresh, young, green leaves were sampled when the reference plant was approximately six months old, and 2 g tissue was flash frozen in liquid nitrogen and ground in a mortar. High molecular weight (HMW) DNA for long read sequencing was extracted using the NucleoBond HMW kit (*NucleoBond HMW DNA*, n.d.) following the protocol except for extending the lysis time to three hours.

Hi-C libraries were built in the Todesco Lab at the University of British Columbia following a detailed Hi-C protocol for plants (Hirabayashi et al., 2025), using 1 g leaf tissue from the reference plant ground with liquid nitrogen. Nuclei were fixed using formaldehyde and isolated. Chromatin digestion was performed using the DpnII restriction enzyme; fragments were then biotinylated and ligated. About 4 µg extracted DNA was used to build an Illumina whole genome sequencing library.

PacBio HiFi libraries were prepared and sequenced by the Centre d'expertise et de services Génome Québec, resulting in 30x coverage. Hi-C libraries were sequenced on the Illumina HiSeqX platform at the Centre d'expertise et de services Génome Québec; 14.2 Gbp of data were produced (18x coverage).

PacBio long reads and Hi-C Illumina short reads were checked for quality using FastQC (Andrews, 2010). Hifiasm was used to make the initial contig assembly with default settings for a phased diploid assembly (Cheng et al., 2021). The contig level assembly was polished with Longstitch. This took the same long reads and remapped them to the contig assembly to polish and run initial scaffolding (Coombe et al., 2021). Illumina Hi-C reads were mapped to the assembly with nf-core HiC-Pro (Servant et al., 2015). The assembly was scaffolded with Juicer (Durand, Robinson, et al., 2016; Durand, Shamim, et al., 2016) and 3D DNA (Dudchenko et al., 2017).

Assembly statistics for both haplotypes of the phased assembly were checked using the assembly-stats tool (Hunt et al., 2021). BUSCO (v5.1.2) (Manni et al., 2021) was run against the BUSCOv5 lineage eudicots_odb10 database (Manni et al., 2021) to assess the completeness of the assembly.

C: Flow cytometry protocol

Before starting, for every sample–standard combination, prepare a filter by placing two layers of Miracloth between plastic pieces, cut the ends off the 1 mL pipette tips, and label the 2 mL tubes. Seeds to grow the standard plants (Table C.1) were received in November 2021 from Prof. Ing. Jaroslav Doležel, DrSc. [(http://olomouc.ueb.cas.cz](%20(http://olomouc.ueb.cas.cz)).

**Table C.1: Standard plants for flow cytometry and their 2C DNA content.**

| **Standard species** | **2C DNA content** |
| --- | --- |
| *Raphanus sativus* | 1.11 pg |
| *Solanum lycopersicum* | 1.96 pg |
| *Glycine max* | 2.50 pg |
| *Zea mays* | 5.43 pg |
| *Pisum sativum* | 9.09 pg |
| *Secale cereale* | 16.19 pg |
| *Vicia faba* | 26.90 pg |

To prepare the buffer, add the reagents in Table C.2 to approximately 150 mL of distilled water. Adjust the volume to 250 mL and the pH to 7. Filter the buffer to remove contaminants, then add the β-mercaptoethanol.

Table C.1: Nuclei extraction and wash buffer was slightly modified from de Laat's buffer (de Laat & Blaas, 1984) and based off of the Rieseberg Lab blog: <https://www.zoology.ubc.ca/~rieseberg/RiesebergResources/dna-ploidy-and-genome-size-estimation-using-flow-cytometry/>

| **For 250mL buffer** | **Amount** |
| --- | --- |
| 0.894g | 15 mM HEPES |
| 0.5mL of 0.5M | 1mM EDTA |
| 0.5mL | 0.2 % (v/v) Triton X-100 |
| 1.49g | 80 mM KCl |
| 0.292g | 20 mM NaCl |
| 25.7g | 300 mM sucrose |
| 12.5mL | 0.5 mM spermine |
| 0.262mL – add after filtering | 15 mM β-mercaptoethanol |
| 2.5g | 0.25 mM PVP-40 |

**Lab Procedure**

1. Measure 30–40 mg of leaf tissue into a Petri dish for each standard–standard, standard–sample, and sample–sample combination. Keep the tissue moist and refrigerated while it waits to be cut.
2. On a chilled aluminum block, rapidly chop the tissue for approximately 1 minute in 1 mL of cold buffer.
3. Using the prepared cut-tip pipette, transfer the buffer and tissue onto the double-layer Miracloth filter positioned over the labeled 2 mL tube. Allow all the liquid to pass through.
4. Centrifuge at room temperature for 5 minutes at 1,200 rpm (slow speed). A white or sometimes green pellet should form. Carefully pipette off the supernatant; the pellet may not be firmly attached to the tube wall.
5. Resuspend the pellet in 1 mL of buffer and repeat the centrifugation and supernatant-removal step.
6. Fix the nuclei overnight at 4°C in ethanol–acetic acid (3:1, v/v). Mix well by inversion.
7. Repeat the centrifugation and supernatant-removal step. If the pellet does not adhere to the tube wall, centrifuge for an additional minute at 1,500 rpm.
8. Add 200 mL of buffer.
9. Add 4 µL of 20 mg/mL RNase A and incubate for approximately 30 minutes at room temperature.
10. Immediately before taking the sample to the flow cytometer, add 20 µL of propidium iodide to the 200 mL of buffer. Adding more dye may broaden the peaks.

**Data analysis**

From the fluorescence peaks, we can calculate the genome size:
Sample_2C _pg = (Standard_2C_pg * mean_sample_peak)/ mean_standard_peak

And then convert DNA content in picograms to base pairs using:
Diploid_bp = 0.978 *10^9^ * Sample_2C_pg

D: TimeTree

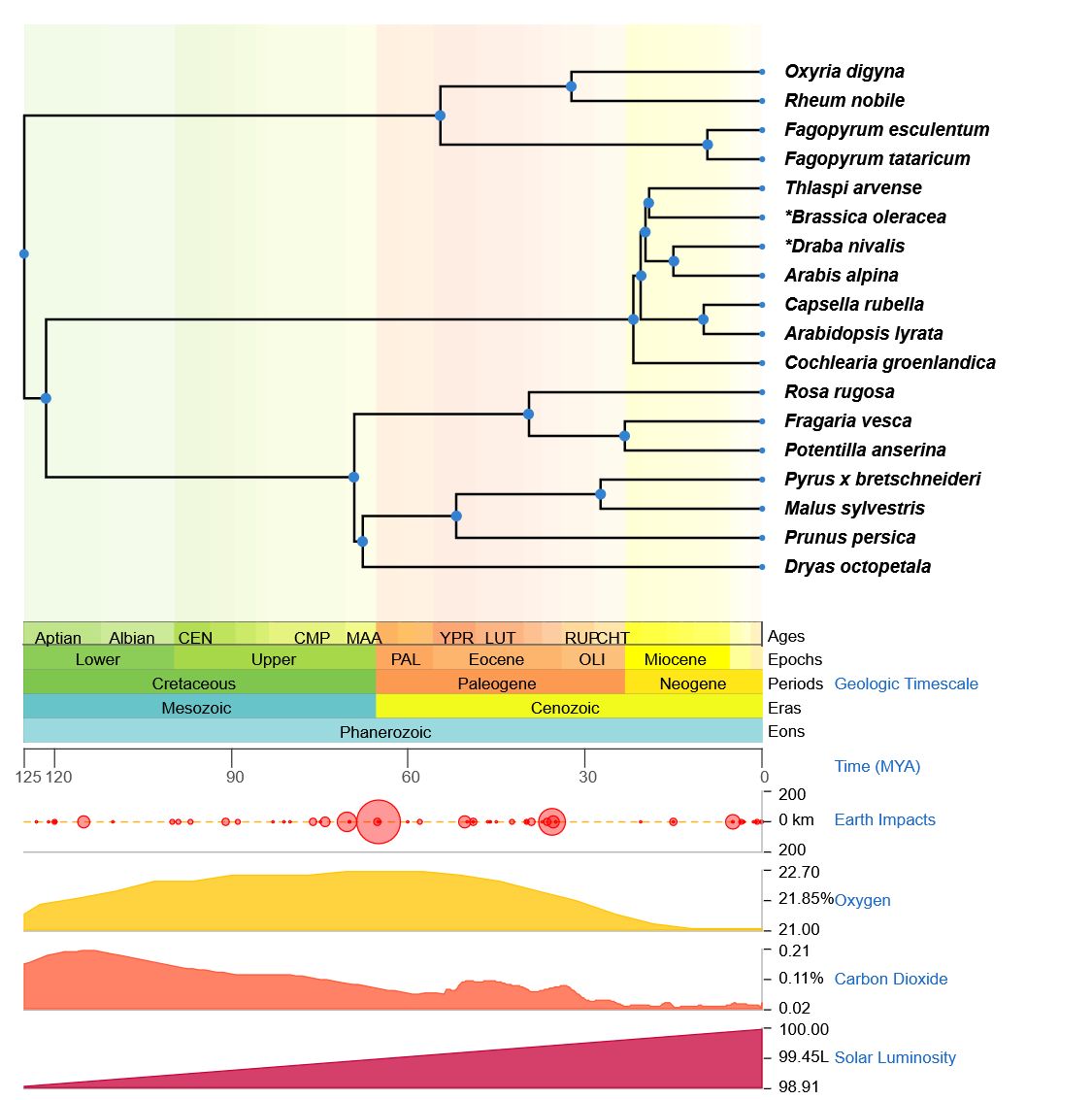
Figure D.1: TimeTree plot from (Kumar et al., 2017).

E: CAFÉ Expanded and contracted gene ontology results

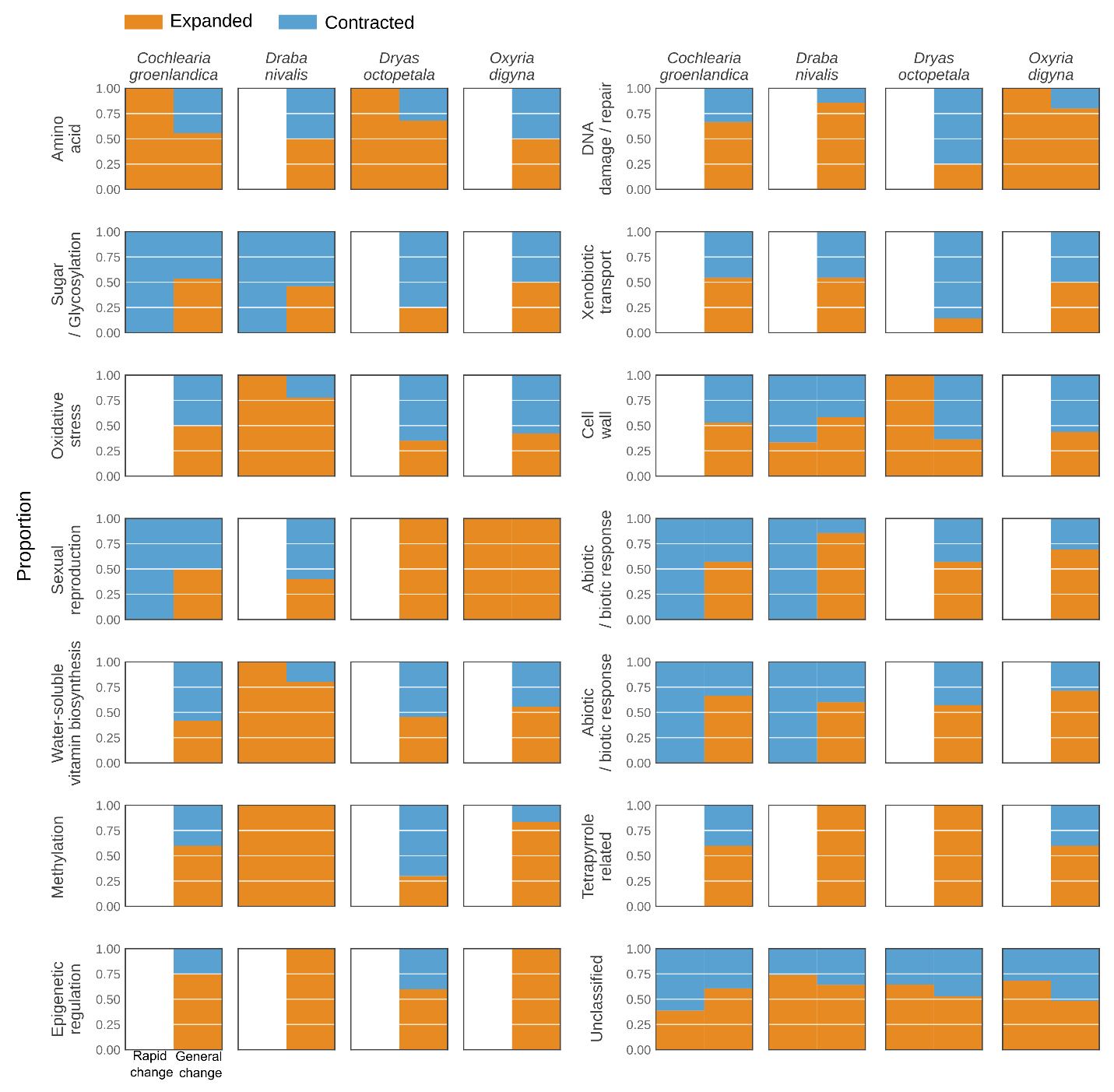

Figure E1: Fraction of changing orthogroups that are contracting versus expanding for each functional category.
